# GABA_B_ receptor-mediated potentiation of ventral medial habenula glutamatergic transmission in GABAergic and glutamatergic interpeduncular nucleus neurons

**DOI:** 10.1101/2025.01.03.631193

**Authors:** Hannah E. Stinson, Ipe Ninan

## Abstract

The medial habenula (MHb)-interpeduncular nucleus (IPN) pathway plays an important role in information transferring between the forebrain and the midbrain. The MHb-IPN pathway has been implicated in the regulation of fear behavior and nicotine addiction. The synapses between the ventral MHb and the IPN show a unique property, i.e., an enhancement of synaptic transmission upon activation of GABA_B_ receptors. This GABA_B_ receptor-mediated potentiation of ventral MHb-IPN synaptic transmission has been implicated in regulating fear memory. Although IPN is known to contain parvalbumin (PV) and somatostatin (SST) GABAergic neurons and vesicular glutamate transporter 3 (VGLUT3)-expressing neurons, it is unknown how GABA_B_ receptor activation affects ventral MHb-mediated glutamatergic transmission onto these three subtypes of IPN neurons. Our studies show robust glutamatergic connectivity from ventral MHb to PV and SST neurons in the IPN, while the ventral MHb-mediated glutamatergic transmission in IPN VGLUT3 neurons is weak. Although activation of GABA_B_ receptors produces a robust potentiation of ventral MHb-mediated glutamatergic transmission in PV neurons, we observed a modest effect in IPN SST neurons. Despite the diminished basal synaptic transmission between ventral MHb and IPN VGLUT3 neurons, activation of GABA_B_ receptors causes transient conversion of non-responding ventral MHb synapses into active synapses in some IPN VGLUT3 neurons. Thus, our results show strong ventral MHb connectivity to GABAergic IPN neurons compared to VGLUT3-expressing IPN neurons. Furthermore, GABA_B_ receptor activation produces a differential effect on ventral MHb-mediated glutamatergic transmission onto PV, SST, and VGLUT3 neurons in the IPN.

## Introduction

The dorsal diencephalic conduction system comprised of the medial habenula (MHb)-interpeduncular nucleus (IPN) pathway is critical for information processing between the forebrain and the midbrain (Sutherland, 1982). The MHb-IPN pathway plays a crucial role in nicotine addiction and fear suppression (Antolin-Fontes et al., 2015; Koppensteiner et al., 2017; Melani et al., 2019; Soria-Gomez et al., 2015; Zhang et al., 2016). The MHb contains two major subdivisions: 1) the dorsal MHb and 2) the ventral MHb (Andres et al., 1999; Contestabile et al., 1987; Kobayashi et al., 2013; Quina et al., 2017; Yamaguchi et al., 2013). While the substance P-expressing dorsal MHb neurons project to the lateral IPN, the choline acetyltransferase (ChAT)-expressing ventral MHb neurons send axons to the rostral and central IPN (Frahm et al., 2015; Melani et al., 2019). A notable property of MHb-IPN synapses is the involvement of multiple neurotransmitters and neuropeptides in this pathway (Frahm et al., 2015; Melani et al., 2019; Zhang et al., 2016). Our studies have shown that substance P enhances activity-dependent plasticity at dorsal MHb-lateral IPN synapses through endocannabinoid CB1 receptor-mediated suppression of GABA_B_ receptor activity (Melani et al., 2019). Furthermore, activation of the NK1 receptor, the primary receptor for substance P, in the IPN is involved in fear extinction (Melani et al., 2019). The ventral MHb neurons release glutamate and acetylcholine onto the rostral and central IPN (Zhang et al., 2016). A remarkable feature of ventral MHb-mediated transmission into the IPN is its role in suppressing fear behavior (Zhang et al., 2016) (Melani et al., 2019). Another unique characteristic of ventral MHb-IPN synapses is the excitatory role of GABA_B_ receptors in enhancing synaptic transmission, which is mediated by an augmentation of Ca^2+^ entry through the calcium channel, Cav2.3 (Bhandari et al., 2021; Koppensteiner et al., 2024; Koppensteiner et al., 2017; Zhang et al., 2016). This effect is consistent with the robust pre-synaptic expression of Cav2.3 channels at MHb-IPN synapses (Bhandari et al., 2021; Parajuli et al., 2012). A recent study shows that K+-channel tetramerization domain-containing proteins regulate synaptic strength at the MHb-IPN synapses independent of GABA_B_ receptors (Bhandari et al., 2021). Furthermore, activation of GABA_B_ receptors at the ventral MHb-IPN synapses enhances fear extinction (Zhang et al., 2016). Our earlier studies show that activity-dependent plasticity at the MHb-IPN glutamatergic synapses depends upon a post-synaptic calcium-permeable AMPA receptor-mediated GABA release and retrograde activation of GABA_B_ receptors on MHb terminals (Koppensteiner et al., 2017). Using multiple experimental approaches, it was recently demonstrated that GABA_B_ receptor activation of MHb terminals induces an activity-dependent transition from a tonic to a phasic neurotransmitter release at MHb-IPN synapses (Koppensteiner et al., 2024).

Although IPN is believed to be comprised of primarily GABAergic neurons, a characterization of GABAergic neurons present in the IPN is lacking (Quina et al., 2017). In addition to the GABAergic neurons, studies suggest the presence of glutamatergic neurons expressing vesicular glutamate transporter 3 (VGLUT3) in the IPN (Herzog et al., 2004; Quina et al., 2017). However, it is unknown whether the robust effect of GABA_B_ receptor activation on glutamate release at the ventral MHb-IPN synapses is specific to any subtypes of IPN neurons. Therefore, we asked whether GABA_B_ receptor activation affects ventral MHb-mediated glutamatergic transmission in two subtypes of GABAergic neurons, i.e., somatostatin (SST) and parvalbumin (PV) neurons and VGLUT3-expressing neurons in the IPN. Our results show that activation of GABA_B_ receptors produces a robust potentiation of ventral MHb-mediated glutamatergic transmission in PV neurons. In contrast, SST neurons show a modest potentiation of ventral MHb-mediated glutamatergic transmission. Furthermore, ventral MHb-mediated glutamatergic transmission in IPN VGLUT3 neurons is significantly weaker than in IPN GABAergic neurons. However, activation of GABA_B_ receptors causes a transient conversion of non-responding ventral MHb synapses into active synapses in some IPN VGLUT3 neurons.

## Results

Earlier studies have demonstrated that GABA_B_ receptor activation by perfusion of an agonist, baclofen (1 µM), induced a multifold increase in ventral MHb-mediated glutamatergic transmission in the central IPN neurons (Bhandari et al., 2021; Koppensteiner et al., 2024; Koppensteiner et al., 2017; Zhang et al., 2016). Consistently, our whole-cell recording experiments in central IPN neurons showed baclofen-induced robust potentiation of light-evoked glutamatergic transmission at ventral MHb-central IPN synapses from brain slices of ChAT ChR2-EYFP mice (Figure 1). The ChAT-positive ventral MHb terminals release both acetylcholine and glutamate in the IPN (Zhang et al., 2016). The ventral MHb-mediated glutamatergic currents were recorded as described before (Koppensteiner et al., 2017). The amplitudes of excitatory post-synaptic currents were 34.47±7.25pA and 43.65±7.2pA, respectively, for female and male groups. The baclofen perfusion showed a 6-7-fold increase in the EPSC amplitude in central IPN neurons (Figure 1). We did not observe an effect of sex on baclofen-induced enhancement of glutamatergic transmission at ventral MHb-IPN synapses.

**Figure 1.**
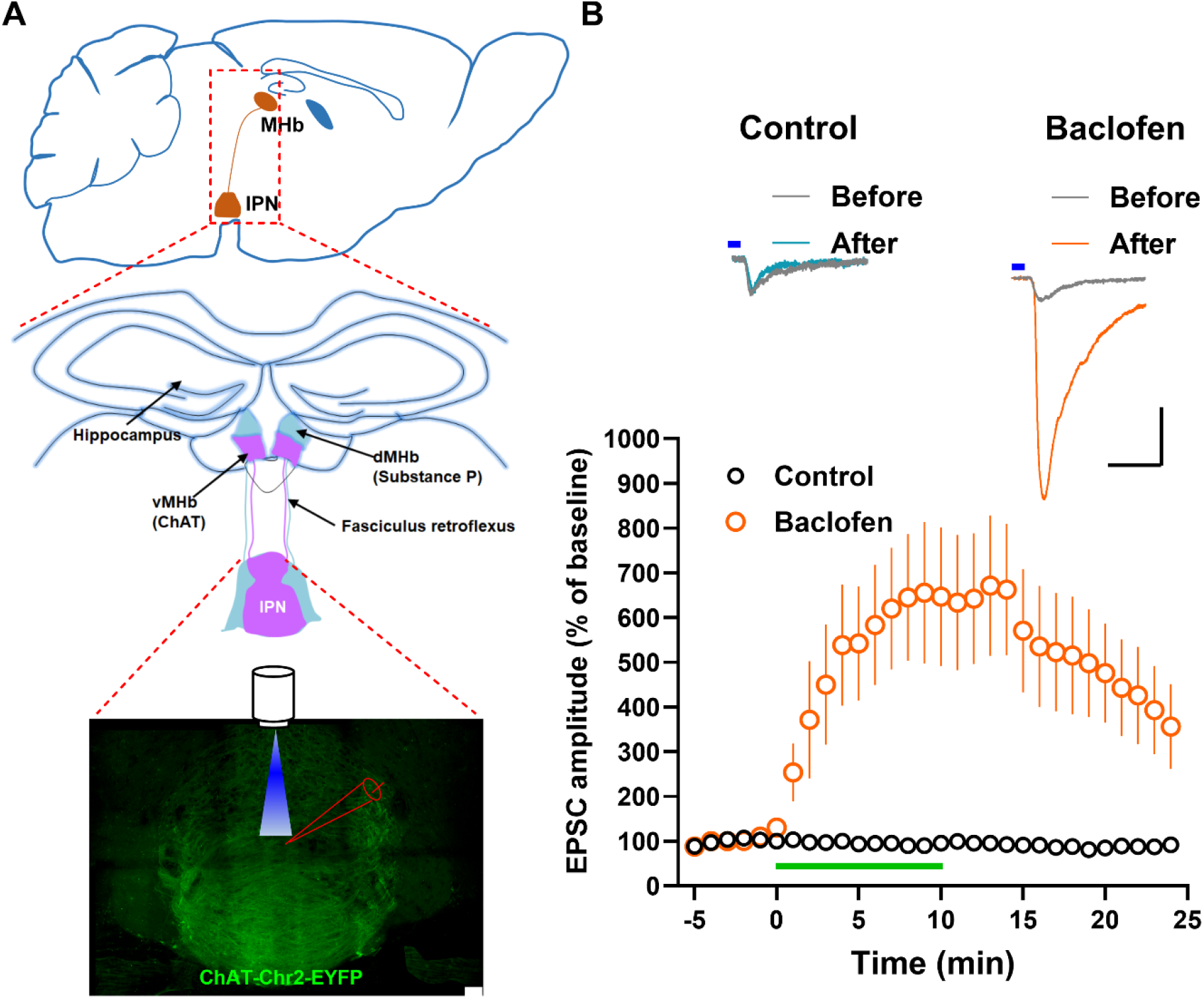
Effect of GABA_B_ receptor activation on ventral MHb-IPN glutamatergic transmission. **A**) A schematic of the MHb-IPN pathway. The bottom panel shows an image of IPN from ChAT-ChR2-EYFP mice. Scale 10 μm. **B**) Effect of baclofen (1 µM), a GABA_B_ agonist, on light-evoked EPSCs in IPN neurons from ChAT-ChR2-EYFP mice. Baclofen group (8 neurons/5 male mice, 10 neurons/5 female mice), control group (13 neurons/4 mice). Baclofen perfusion for 10 mins showed a statistically significant enhancement of EPSC amplitude (F1,29=10.9, P=0.003). We did not observe an effect of sex on baclofen-induced enhancement of EPSC amplitude (P=0.14). The upper panel shows example traces. Scale 100pA/20ms. The blue line indicates light application.

Next, we examined whether baclofen enhances ventral MHb-mediated glutamatergic transmission onto SST-expressing neurons in the IPN. SST-expressing neurons are primarily localized to the rostral part of the IPN (Figure 2). To target SST neurons in the IPN, first, we generated SST-tdTomato mice by crossing Sst-IRES-Cre mice with Ai14(RCL-tdT)-D mice. Then, SST-tdTomato mice were crossed to ChAT-ChR2-EYFP mice to generate ChAT-ChR2-EYFP/SST-tdTomato mice. We undertook whole-cell recording in tdTomato-expressing SST neurons in the rostral IPN. The amplitudes of excitatory post-synaptic currents were 28.32±6.4pA and 31.7±4.9pA, respectively, for female and male groups. Baclofen perfusion for 10 min showed a modest increase in light-evoked EPSC amplitude in the SST neurons (Figure 2). We did not observe an effect of sex on baclofen-induced enhancement of glutamatergic transmission in SST neurons.

**Figure 2.**
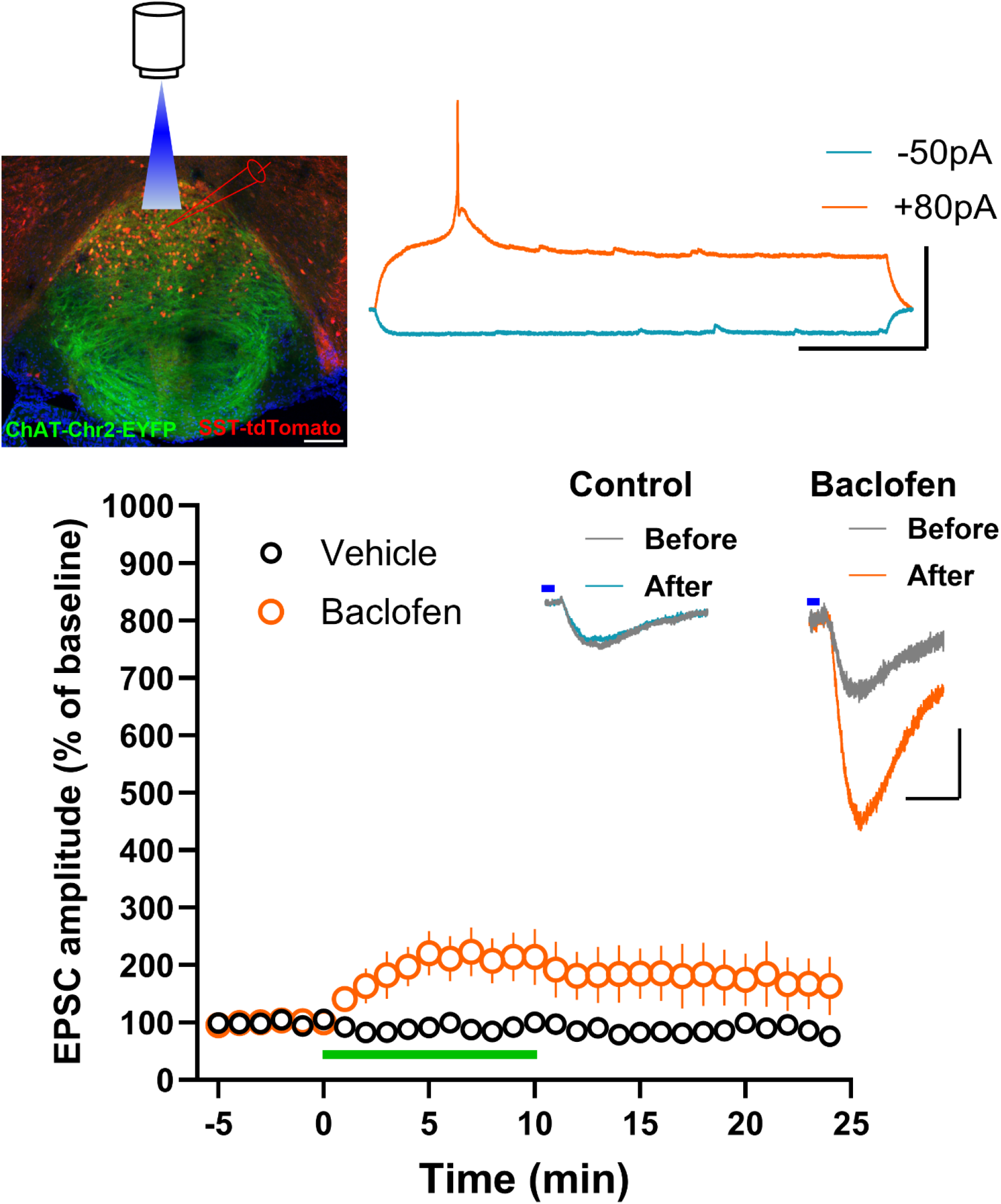
Effect of GABA_B_ receptor activation on ventral MHb-IPN (SST) neuron glutamatergic transmission. The graph shows the effect of baclofen on light-evoked EPSCs in SST-expressing IPN neurons from ChAT-ChR2-EYFP/SST-tdTomato mice. Baclofen group (7 neurons/5 male mice, 9 neurons/5 female mice), control group (8 neurons/4 mice). Baclofen perfusion for 10 mins showed a statistically significant enhancement of EPSC amplitude (F1,22=4.5, P=0.045). We did not observe an effect of sex on baclofen-induced enhancement of EPSC amplitude (P=0.18). The inset shows example traces. Scale 100pA/20ms. The blue line indicates light application. The upper left panel shows an image of IPN from ChAT-ChR2-EYFP/SST-tdTomato mice. Scale 50 μm. The upper right panel shows example traces for voltage changes in response to current injection in IPN (SST) neurons. Scale 50mV/250ms.

To study the effect of baclofen on ventral MHb-mediated glutamatergic transmission in PV-expressing IPN neurons, first, we generated PV-tdTomato mice by crossing B6 PValb-IRES-Cre with Ai14(RCL-tdT)-D mice. Then, PV-tdTomato mice were crossed with ChAT-ChR2-EYFP mice to generate ChAT-ChR2-EYFP/PV-tdTomato mice. Consistent with our previous study, we found the presence of tdTomato-expressing PV neurons in rostral, central, and lateral IPN (Melani et al., 2019). We recorded tdTomato-expressing PV neurons in the central IPN (Figure 3). The amplitudes of excitatory post-synaptic currents were 33.73±11.5pA and 27.65±4.96pA, respectively, for female and male groups. Baclofen showed a robust 6-7 fold potentiation of light-evoked EPSCs in PV neurons in the central IPN (Figure 3). Similar to SST neurons, we did not observe an effect of sex on baclofen-induced potentiation of ventral MHb-mediated glutamatergic transmission in PV-expressing IPN neurons.

**Figure 3.**
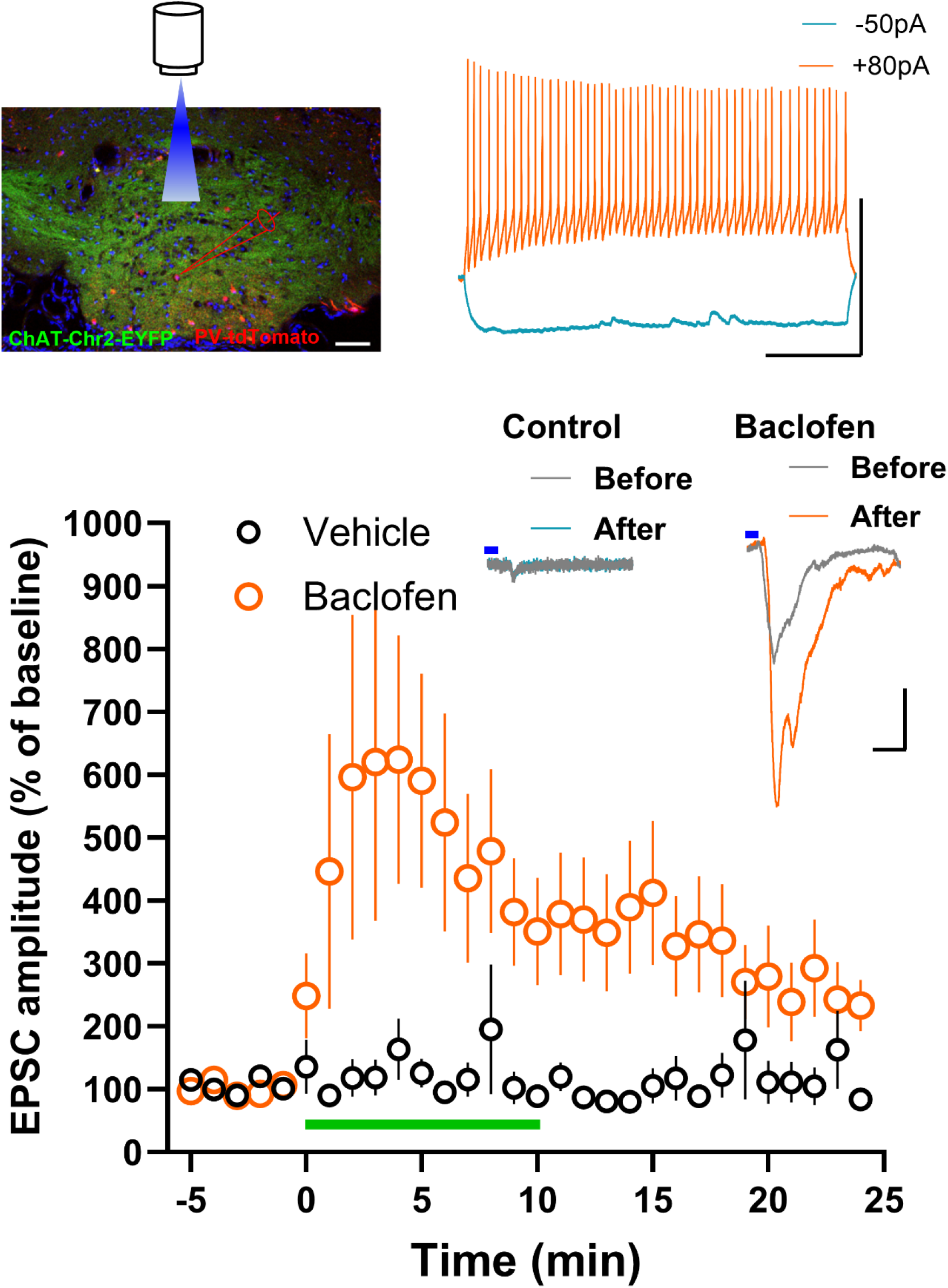
Effect of GABAB receptor activation on ventral MHb-IPN (PV) neuron glutamatergic transmission. The graph shows the effect of baclofen on light-evoked EPSCs in PV-expressing IPN neurons from ChAT-ChR2-EYFP/PV-tdTomato mice. Baclofen group (11 neurons/7 male mice, 12 neurons/6 female mice), control group (9 neurons/7 mice). Baclofen perfusion for 10 mins showed a statistically significant enhancement of EPSC amplitude (F1,30=8.2, P=0.007). We did not observe an effect of sex on baclofen-induced enhancement of EPSC amplitude (P=0.46). The inset shows example traces. Scale 100pA/20ms. The blue line indicates light application. The upper left panel shows an image of IPN from ChAT-ChR2-EYFP/PV-tdTomato mice. Scale 50 μm. The upper right panel shows example traces for voltage changes in response to current injection. Scale 50mV/250ms.

The IPN contains VGLUT3-expressing neurons. To determine whether GABA_B_ receptor activation affects ventral MHb-mediated glutamatergic transmission in IPN VGLUT3 neurons, we generated VGLUT3-tdTomato mice by crossing VGLUT3 Cre mice with Ai14(RCL-tdT)-D mice. Then, the VGLUT3-tdTomato mice were crossed with ChAT-ChR2-EYFP mice to generate ChAT-ChR2-EYFP/VGLUT3-tdTomato mice. Unlike the SST and PV neurons, only 1 out of 24 recorded neurons showed light-evoked ventral MHb-mediated glutamatergic current in central IPN VGLUT3 neurons. While the perfusion of baclofen produced a modest increase in EPSC amplitude in this neuron, 5 out of 9 neurons that showed no response initially were modestly and transiently potentiated by baclofen perfusion (Figure 4).

**Figure 4.**
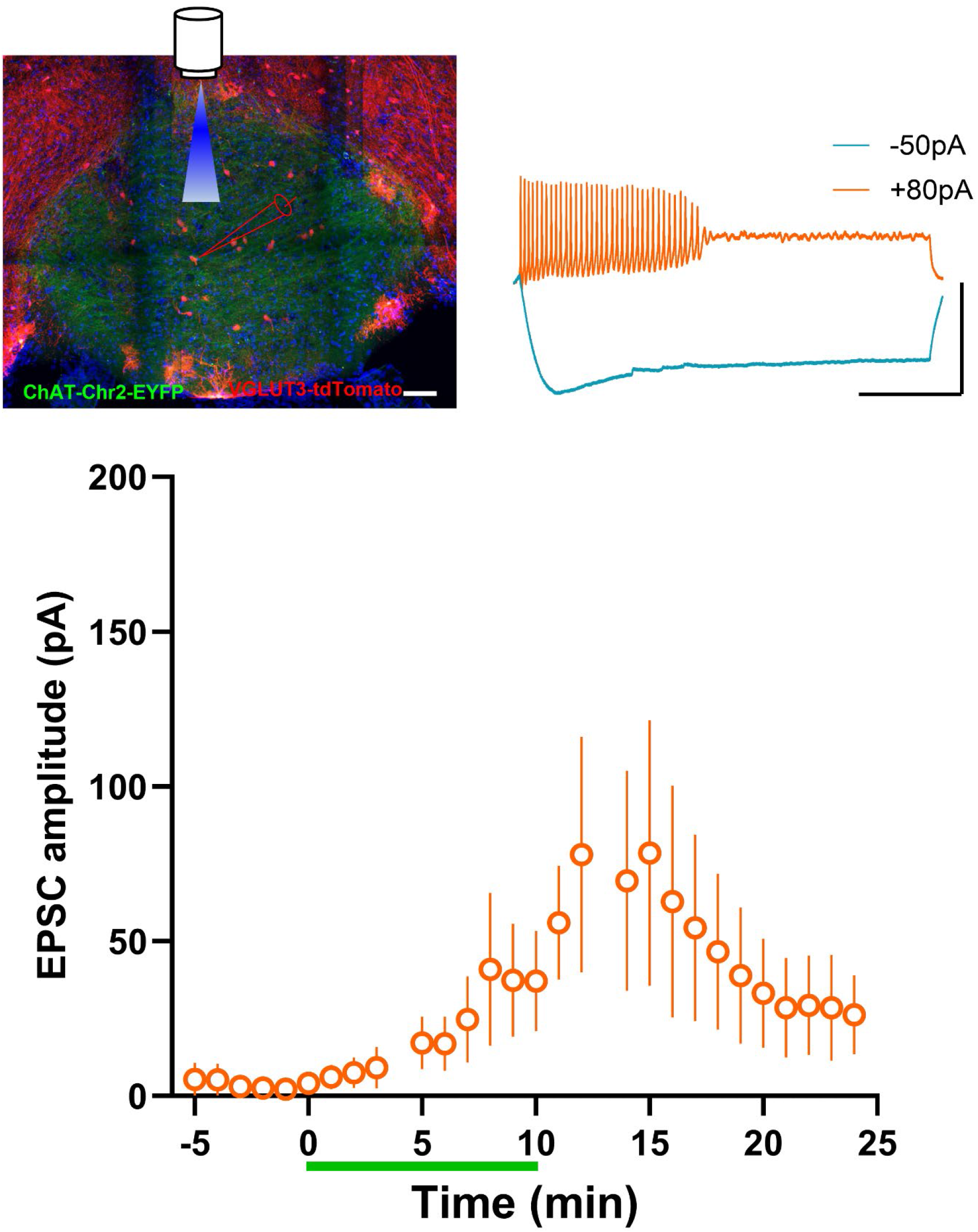
Effect of GABA_B_ receptor activation on ventral MHb-IPN (VGLUT3) neuron glutamatergic transmission. Only 1 out of 24 recorded VGLUT3 neurons showed EPSC in response to light stimulation. Baclofen showed a modest and transient increase in EPSC amplitude in this responding neuron. When baclofen was tested on 9 non-responding neurons, 5 neurons showed a modest and transient increase in EPSC amplitude after baclofen perfusion. The graph shows the effect of baclofen on light-evoked EPSCs in VGLUT3-expressing IPN neurons (1 responding neuron +5 non-responding neurons) from ChAT-ChR2-EYFP/VGLUT3-tdTomato mice. This effect was not statistically significant (F1,4=3.94, P=0.11). The upper left panel shows an image of IPN from ChAT-ChR2-EYFP/VGLUT3-tdTomato mice. Scale 50 μm. The upper right panel shows example traces for voltage changes in response to current injection. Scale 50mV/250ms.

These results show strong ventral MHb connectivity to GABAergic IPN neurons compared to VGLUT3-expressing IPN neurons. In addition, GABA_B_ receptor activation produces a differential effect on ventral MHb-mediated glutamatergic transmission onto different subtypes of IPN neurons.

## Discussion

Our study shows the strength of ventral MHb glutamatergic synapses onto three different subtypes of IPN neurons. While both PV and SST IPN neurons showed robust synaptic inputs from ventral MHb neurons, the ventral MHb input to the VGLUT3 neurons in the IPN is weak, suggesting a preferential ventral MHb-mediated synaptic transmission onto GABAergic neurons compared to VGLUT3 expressing neurons in the IPN. Activation of GABA_B_ receptors produces a robust potentiation of glutamatergic transmission at the ventral MHb-IPN PV neuron synapses. Although we observed GABA_B_ receptor-mediated enhancement of glutamatergic transmission at the ventral MHb-IPN SST neuron synapses, the magnitude of the effect of GABA_B_ receptor activation was lower in SST neurons than in PV neurons. Although the ventral MHb-IPN VGLUT3 neuron synaptic transmission is significantly weak, activation of GABA_B_ receptors transiently converted non-responding ventral MHb synapses to active synapses in VGLUT3 neurons. These results show differential effects of GABA_B_ receptor activation on ventral MHb-mediated glutamatergic transmission onto PV, SST, and VGLUT3 neurons in the IPN.

Unlike the well-known inhibitory role of GABA_B_ receptors, activation of GABA_B_ receptors at the ventral MHb-IPN synapses potentiates glutamatergic and cholinergic transmission (Bhandari et al., 2021; Koppensteiner et al., 2024; Koppensteiner et al., 2017; Zhang et al., 2016). In contrast to ventral MHb-IPN synapses, activation of GABA_B_ receptors suppresses dorsal MHb-mediated transmission onto lateral IPN neurons (Melani et al., 2019). Furthermore, activation of GABA_B_ receptors at the ventral MHb-IPN synapses enhances fear extinction (Zhang et al., 2016). Given the effect of GABA_B_ receptor activation on ventral MHb-mediated glutamatergic transmission onto three subtypes of IPN neurons, it is highly likely that GABA_B_ receptor activation modulates IPN-mediated regulation of downstream targets, which include the median raphe nucleus, central gray, and nucleus incertus (Hsu et al., 2013; Quina et al., 2017). However, little is known about where these three subtypes of IPN neurons project and whether they mediate specific effects on behavior. Recent studies show the role of IPN GABAergic neurons in anxiolytic stress-coping mechanisms (Klenowski et al., 2023). Specifically, IPN SST neurons play a crucial role in stress-induced increase in activity in the IPN and stress-induced coping mechanisms (Klenowski et al., 2023). IPN SST neurons also exert a pre-synaptic effect to regulate MHb-mediated glutamate release in the IPN (Zhao-Shea et al., 2013). Given that both stress and nicotine affect SST neuron activity in the IPN, GABA_B_ receptor-mediated enhancement of glutamatergic transmission onto IPN SST neurons might influence stress and nicotine withdrawal symptoms (Klenowski et al., 2023; Zhao-Shea et al., 2013). SST neurons are primarily confined to the rostral IPN, which project to the central gray and nucleus incertus (Quina et al., 2017). The nucleus incertus-projecting IPN neurons are critical for amplifying aversion and promoting avoidant behaviors (Liang et al., 2024). SST neurons, which also express α5 nicotinic receptors, project to the medial raphe nucleus (Hsu et al., 2013). Therefore, the effect of GABA_B_ receptor activation on IPN SST neurons is expected to modulate the median raphe nucleus, which projects to several forebrain areas (Collins et al., 2023; Fortin-Houde et al., 2023; Stinson and Ninan, 2024; Szonyi et al., 2016). Since IPN projections are primarily descending but not ascending in nature, the effect of GABA_B_ receptor activation on IPN neurons is expected to modulate forebrain structures through these descending pathways (Quina et al., 2017). Also, IPN neurons project to the laterodorsal tegmental nucleus (Quina et al., 2017). Thus, the robust and differential effects of GABA_B_ receptor activation on ventral MHb-mediated glutamatergic transmission onto IPN neurons are expected to affect behaviors pertinent to aversion, reward, and addiction. Given the lack of knowledge on the specific IPN neuron subtypes that project to other brain areas, future studies are needed to understand how subtypes of IPN neurons modulate other brain structures. In addition to PV, SST, and VGLUT3 IPN neurons, IPN shows the presence of VGLUT2 glutamatergic neurons, which project to the laterodorsal tegmental nucleus (Quina et al., 2017). Therefore, future studies are needed to understand how PV-, VGLUT3- and VGLUT2-expressing IPN neurons modulate other brain areas and determine whether and how GABA_B_ receptor activation in the IPN modulate those behaviors.

## Materials and Methods

### Mice

2-3 months old female and male mice were used. *Sst*^*tm2*.*1(cre)Zjh*^/J (Stock number: 013044, the Jackson Laboratories), B6.129P2-*Pvalb*^*tm1(cre)Arbr*^/J (Stock number: 017320, the Jackson Laboratories), and B6;129S-*Slc17a8*^*tm1*.*1(cre)Hze*^/J (Stock number: 028534, the Jackson Laboratories) mice were crossed with B6.Cg-*Gt(ROSA)26Sor*^*tm14(CAG-tdTomato)Hze*^/J (Stock number: 007914, the Jackson Laboratories) to generate PV-tdTomato, SST-tdTomato and VGLUT3-tdTomato mice. B6.Cg-Tg(Chat-COP4*H134R/EYFP,Slc18a3)6Gfng/J (ChAT-ChR2-EYFP, Stock number: 014546, the Jackson Laboratories) mice were used for crossing with PV-tdTomato, SST-tdTomato and VGLUT3-tdTomato mice. Mice were maintained on a 12-hour light-dark cycle at 23°C with access to food and water ad libitum. The Institutional Animal Care and Use Committee of the University of Toledo approved all the procedures.

### Electrophysiology

Mice underwent intracardiac perfusion with ice-cold oxygenated artificial cerebrospinal fluid (ACSF) containing (in nM): NaCl (118), glucose (10), KCl (2.5), NaH_2_PO_4_ (1), CaCl_2_ (1) and MgSO_4_ (1.5) (325 mOsm, pH 7.4) under pentobarbital (120mg//kg) anesthesia. IPN slices (250 µm) were prepared on a vibratome after the quick removal of the brain. Brain slices were collected in oxygenated ACSF at 35°C and allowed to cool to room temperature during incubation for I hr. Brain slices were transferred to a recording chamber superfused with the ACSF containing 2.5 mM CaCl_2_ at 32°C flowing at 2 ml/min rate on an upright microscope (Zeiss Examiner D1). The IPN was located using a 4X objective. Recorded IPN neurons were visualized using a 40X water immersion objective and video-enhanced differential interference contrast microscopy. tdTomato-expressing IPN neurons were identified using fluorescence microscopy. IPN neurons were recorded using glass electrodes of 3–5 MΩ resistance filled with an internal solution containing (in mM): K-gluconate (130), KCl (10), MgCl_2_ (5), MgATP (5), NaGTP (0.2), HEPES (5), pH adjusted to 7.4 with KOH. Ventral MHb-mediated currents in IPN neurons were recorded at -60mV by light stimulation (5 ms, 100% light intensity) as described before (Koppensteiner et al., 2017). Blue light (470 nm) was emitted through the 40X water immersion objective from a Lumen 1600-LED (Prior). To study the effect of GABA_B_ receptor activation on ventral MHb-mediated glutamatergic transmission in IPN neurons, baclofen (1 µM) was perfused in ACSF for 10 min. Electrophysiological recordings were rejected when series resistance or holding current changed by 10% or more.

We used Clampfit 10.7 (Molecular Devices, CA, USA) to analyze synaptic responses. Statistical analyses were performed using IBM SPSS statistics (version 27) or GraphPad Prism (version 9) software. A two-way repeated measures ANOVA was used for analyzing EPSC amplitude.

### Confocal microscopy

Mice were intracardially perfused with 4% PFA in PBS under pentobarbital (120mg/kg) anesthesia. Brains were collected in 4% PFA and stored at 4°C overnight before being placed in 15% sucrose for 24 hr, followed by 30% sucrose for 24 hr at 4°C. Brains were frozen on dry ice, and 45 µm cryosections of the brain slices were made. Brain slices were mounted on Superfrost Plus microscope slides and coversliped with Fluoromount-G with DAPI (Southern Biotech). Confocal images of the IPN were taken using a TCS SP5 multiphoton laser-scanning confocal microscope (Leica Microsystems, Buffalo Grove, IL) with a 20X objective lens.

